# AncestryHub: A web server for local ancestry analysis

**DOI:** 10.1101/2025.01.02.630692

**Authors:** Sukun Jiang, Yangyang Deng, Shuxin Li, Changsheng Jonathan Liu, Jianjun Luo, Xiaojun Zhu, George D. Song, Kui Zhang, Qing Song, Li Ma

## Abstract

**Summary:** AncestryHub is a web version of local ancestry analysis software. It contains four built-in software and a set of built-in population reference panels. AncestryHub will significantly reduce the efforts and requirements for the users on their personal computational skills and computational hardware environment for ancestry analysis.

**Availability:** https://www.ancestryhub.4dgenome.com

## 1 Introduction

Most of modern populations in the world, if not all, are ancestrally admixed populations, in which different genomic loci of the same individual may have been inherited from different ancestral origins **(Hellenthal et al., 2014; Patterson et al., 2012)**. Accurate high-resolution determination of ancestral origins and stratification of underlying population substructure is not only of broad interest by lay communities but also critical for association studies and genetic studies **(Atkinson et al., 2021; Shriner, Adeyemo, & Rotimi, 2011)**.

Now a series of local ancestry inference software tools have been developed **(Supplementary Note-1)**; however, both academic users and the lay users often encountered some technical issues on computational environments or programming skills when using these software tools. Here, we present a web-based user-friendly platform for local ancestry analysis and visualization, called AncestryHub.

## 2 Implementation

The AncestryHub server contains four built-in software and a set of population reference panels underlying the analytical pipeline.

The built-in reference panels include totally 5,044 human haplotypes from 2522 individuals of 20 human populations from the 1,000 Genomes Project **(Auton et al., 2015)** (**Supplementary Table S3**). The users can also use their own customized reference panels. For the users’ convenience, when the SNP sets of the users’ data files do not match to the SNP sets of the reference panel, AncestryHub will automatically provide an alignment function and pick up the matched SNPs for the local ancestry analysis.

The four built-in software are, 1) a core analytical software for local ancestry analysis, called aMAP (ancestry of Modern Admixed Populations), which can finish a whole genome local-ancestry analysis with a 99.4% accuracy in a very fast computing speed **(Ma et al., 2014)**, 2) a visualization software, called AncestryView, for converting the digital data to a data-driven imag**e (Zhao et al., 2019)**, 3) a phasing software for the users’ convenience to convert the genotype input data to haplotype data, and 4) an intelligent reference switch for selecting the appropriate reference panels.

The web version of this ancestry inference platform has 20 built-in population reference panels for users’ convenience, which belong to 4 super populations, AFR (Africans, 703 individuals), EAS (East Asians, 585 individuals), EUR (Europeans, 633 individuals), and SAS (South Asians, 601 individuals). The data of these populations were downloaded from The 1000 Genomes Project (KGP) **(Auton et al., 2015)**.

The input files are either the genotypes or haplotypes, in either the VCF format. To ensure the fine resolution, we arbitrarily require that the input file should have >=2000 SNPs on each chromosome for local ancestry analysis.

The output contains three files for each individual (**Figure 1 and Figure 2**), (1) an image of local ancestry results of either whole genome or a piece of chromosomal region for direct visualization (**Supplementary Figures S1 and S2**); (2) a digital file of detailed ancestral report along each chromosome for the whole genome for users’ subsequent analysis (**Supplementary Figures S3 and S4**); (3) a summary file in a CSV format (**Supplementary Tables S1 and S2**).

**Figure 1.**
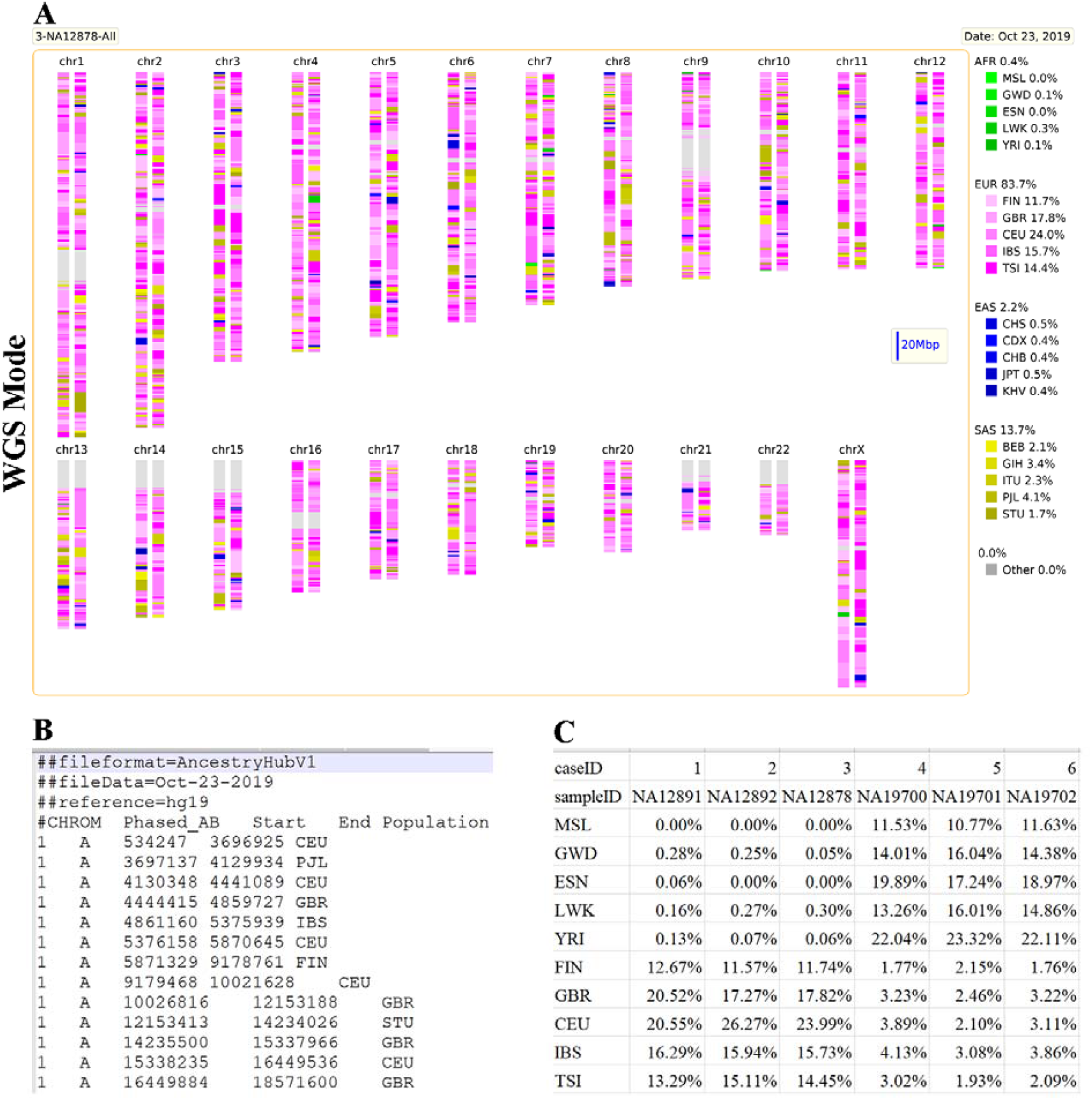
An example of AncestryHub output in WGS mode. The AncestryHub has two modes, which can be used to analyze individual whole-genomes (WGS mode) or single chromosomes or regions (SCA mode). As for WGS mode, it has three output documents for each input, (A) ancestry results in image; (B) ancestry results in digital reports; (C) a summary.

**Figure 2.**
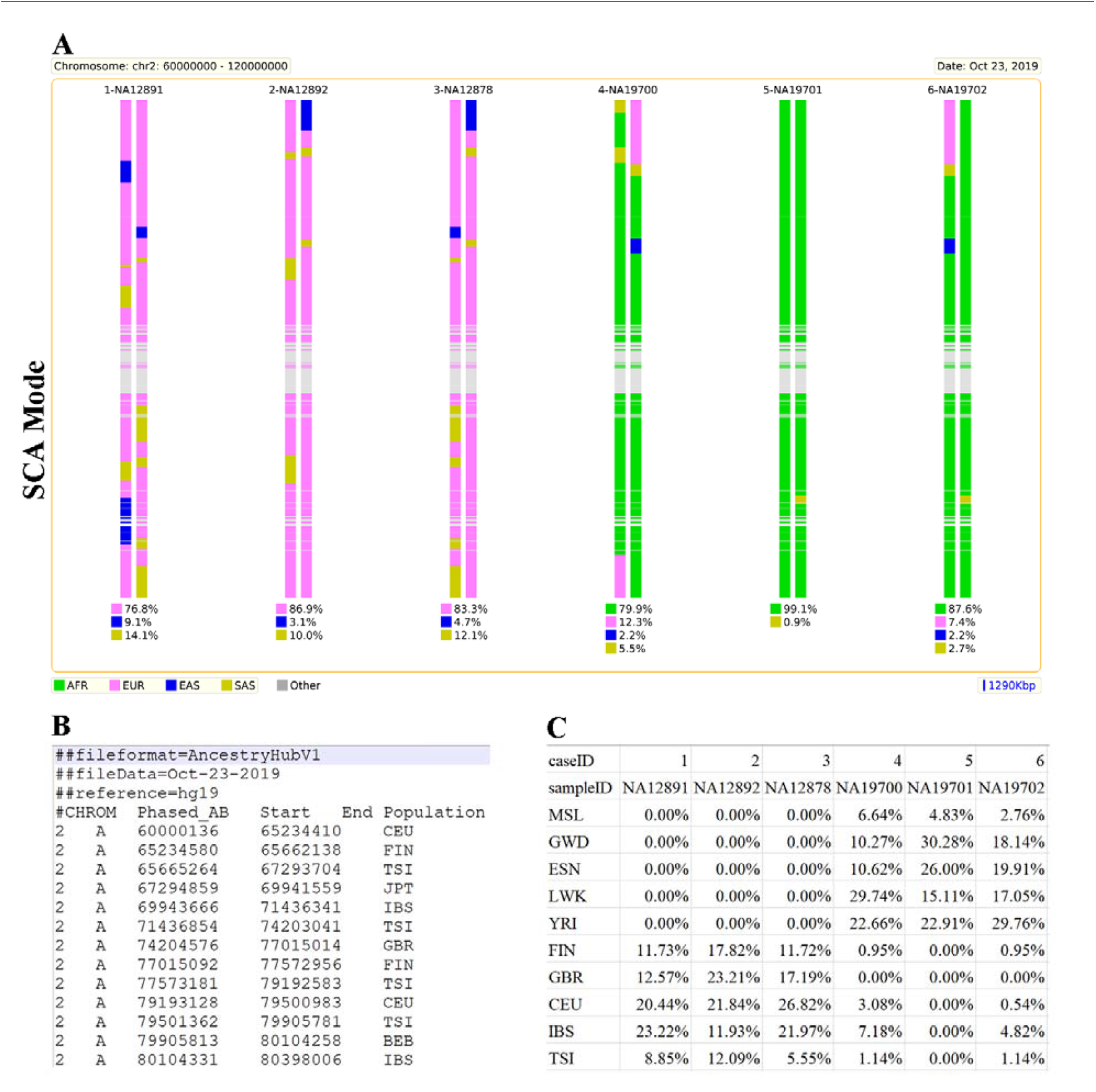
An example of AncestryHub output in SCA mode. As for single chromosomes or regions (SCA mode), the output has three different output documents for each input, (A) ancestry results in image; (B) ancestry results in digital reports; (C) a summary.

The analytical process is composed of the following steps, (i) aligning the SNPs between the input data and references; (ii) phasing; (iii) automatically selecting the appropriate reference panels; (iv) analyzing local ancestry with aMAP; (v) creating the visualization image with AncestryView; (vi) sending a notification email to the users.

## 3 Features and Conclusion

Compared with the aMAP software that we developed previously **(Ma et al., 2014)** and other local ancestry analytical software tools (**Supplementary Note-1**). AncestryHub has the following features:

### High accuracy

The core software aMAP underlying the AncestryHub is featured by high accuracy (99.4%), high speed, and high-resolution.

### High speed

We examined the computing speed of AncestryHub on a regular desktop computer (Intel® Core™ i7-2600K, 32GB RAM). The speed is related to the size of reference panels and the choice of the WGS/SCA modes. It took 45 mins for automatic 20-panal reference preparation for the WGS mode, or 60 seconds for automatic 20-panal 60-Mbp-region reference preparation for the SCA mode.

### Intelligent reference matching

Users do not need to provide the genetic background for this ancestry analysis. AncestryHub will choose the appropriate reference panels for the samples. Briefly, it will carry out three rounds of ancestry analysis, the first round is a pre-analysis for the reference selection, the second round is using the super-population panels, and the third round is using 20 reference panels.

### Customized option between a whole-genome analysis and a target region analysis

Some users may want to zoom-in to a target genomic region where they are specifically interested, so we designed two modes for users, the whole-genome (WGS) mode or specific chromosome area (SCA) mode for the users to choose according to their needs.

### The number of reference panels is large

The aMAP software has an ability to handle a large number of reference panels, this is unique among local ancestry inference software tools. It also has a unique ability to detect “others” when an ancestral region is not covered by the reference panels. It can also distinguish closed related populations, and detect the ancient and small ancestral segments.

We selected two trio-family data as example (CEU-family: NA12891, NA12892, NA12878; ASW-family: NA19700, NA19701, NA19702). Figure S1 and S2 show a result from both WGS mode and SCA mode. By using trio-family data, we can see that child-local-ancestry information inherit from parents-local-ancestry information.

Population stratification is a growing concern in whole-genome association studies and various genetic studies. It can lead to false-positive and false-negative association signals due to systematic differences in allele frequencies between subgroups within an undetected underlying population substructure (**Supplementary Literature**). Several studies have demonstrated that in additional to consider the global ancestry information, the use of local ancestry information is necessary to further reduce false-positive and false-negative association signals due to population stratifications (**Supplementary Literature**). Our software package, AncestryHub, can be used to infer global and local ancestry of a large number of samples efficiently.

## Data availability

The AncestryHub web server is available at https://www.ancestryhub.4dgenome.com

## CRediT author statement

Sukun Jiang: Data curation, Methodology, Software, Investigation, Writing – original draft. Yangyang Deng: Methodology, Visualization. Shuxin Li: Data curation, Methodology. Changsheng Jonathan Liu: Methodology, Software, Investigation. Jianjun Luo: Data curation, Methodology. Xiaojun Zhu: Data curation, Methodology. George D. Song: Investigation, Methodology, Data curation. Kui Zhang: Conceptualization, Methodology. Qing Song: Conceptualization, Software, Methodology, Investigation, Supervision, Funding acquisition, Writing – review & editing. Li Ma: Conceptualization, Software, Methodology, Investigation, Supervision, Funding acquisition, Writing – review & editing. All authors have read and approved the final manuscript.

