## Supplementary material for "AncestryHub: A web server for local ancestry analysis": Supplemenary

Short title: AncestryHub

Sukun Jiang^1,2^, Yangyang Deng^2^, Shuxin Li^1,2^, Changsheng Jonathan Liu^2^, Jianjun Luo^2^, Xiaojun Zhu^2^, George D. Song^2^, Kui Zhang^3^, Qing Song^1,2^, and Li Ma^1,2*^

### 1. Supplementary Note-1. A list of local ancestry inference (LAI) software tools.

### 2. Figure S1. The examples of the image outputs from AncestryHub using the WGS mode.

### 3. Figure S2. The examples of the image outputs from AncestryHub using the SCA mode.

### 4. Figure S3. The examples of the digital outputs from AncestryHub using the WGS mode.

### 5. Figure S4. The examples of the digital outputs from AncestryHub using the SCA mode.

### 6. Table S1. An example of the summary output from AncestryHub using the WGS mode.

### 7. Table S2. An example of the summary output from AncestryHub using the SCA mode.

### 8. Table S3. The population Reference Panels from the 1,000 Genomes Project.

9. Supplementary Literature.

**Supplementary Note-1. A list of local ancestry inference (LAI) software tools**

**aMap**

Ma, Yamin, et al. "Accurate inference of local phased ancestry of modern admixed populations." Scientific reports 4.1 (2014): 5800.

**Flare**

Browning, Sharon R., Ryan K. Waples, and Brian L. Browning. "Fast, accurate local ancestry inference with FLARE." The American Journal of Human Genetics 110.2 (2023): 326-335.

**Ancestral Spectrum Analyzer (ASA)**

Fu H, Shi G. Local Ancestry Inference Based on Population-Specific Single-Nucleotide Polymorphisms-A Study of Admixed Populations in the 1000 Genomes Project. Genes (Basel). 2024 Aug 21;15(8):1099. PMID: 39202458

**Recomb-Mix**

Wei, Yuan, Degui Zhi, and Shaojie Zhang. "Fast and accurate local ancestry inference with Recomb-Mix." bioRxiv (2023): 2023-11.

**SALAI-Net**

Oriol Sabat, Benet, et al. "SALAI-Net: species-agnostic local ancestry inference network." Bioinformatics 38. Supplement_2 (2022): ii27-ii33.

**WINC-pipeline**

Molinaro, Ludovica, et al. "A chromosome-painting-based pipeline to infer local ancestry under limited source availability." Genome Biology and Evolution 13.4 (2021): evab025.

**Gnomix**

Hilmarsson, Helgi, et al. "High resolution ancestry deconvolution for next generation genomic data." bioRxiv (2021): 2021-09.

**MendelImpute**

Chu, Benjamin B., et al. "A fast data-driven method for genotype imputation, phasing and local ancestry inference: MendelImpute. jl." Bioinformatics 37.24 (2021): 4756-4763.

**Ancestryinfer**

Schumer, Molly, Daniel L. Powell, and Russ Corbett‐Detig. "Versatile simulations of admixture and accurate local ancestry inference with mixnmatch and ancestryinfer." Molecular ecology resources 20.4 (2020): 1141-1151.

**LAI-NET**

Montserrat, Daniel Mas, Carlos Bustamante, and Alexander Ioannidis. "Lai-net: Local-ancestry inference with neural networks." ICASSP 2020-2020 IEEE International Conference on Acoustics, Speech and Signal Processing (ICASSP). IEEE, 2020.

**MOSAIC**

Salter-Townshend, Michael, and Simon Myers. "Fine-scale inference of ancestry segments without prior knowledge of admixing groups." Genetics 212.3 (2019): 869-889.

**Ancestry HMM**

Medina, Paloma, et al. "Estimating the timing of multiple admixture pulses during local ancestry inference." Genetics 210.3 (2018): 1089-1107.

**Loter**

Dias-Alves, Thomas, Julien Mairal, and Michael GB Blum. "Loter: a software package to infer local ancestry for a wide range of species." Molecular biology and evolution 35.9 (2018): 2318-2326.

**ELAI**

Guan, Yongtao. "Detecting structure of haplotypes and local ancestry." Genetics 196.3 (2014): 625-642.

**MultiMix**

Churchhouse, Claire, and Jonathan Marchini. "Multiway admixture deconvolution using phased or unphased ancestral panels." Genetic epidemiology 37.1 (2013): 1-12.

**RFMix**

Maples, Brian K., et al. "RFMix: a discriminative modeling approach for rapid and robust local-ancestry inference." The American Journal of Human Genetics 93.2 (2013): 278-288.

**LAMP-LD/LAMP-HAP**

Baran, Yael, et al. "Fast and accurate inference of local ancestry in Latino populations." Bioinformatics 28.10 (2012): 1359-1367.

**SPA**

Yang, Wen-Yun, et al. "A model-based approach for analysis of spatial structure in genetic data." Nature genetics 44.6 (2012): 725-731.

**SupportMix**

Omberg, Larsson, et al. "Inferring genome-wide patterns of admixture in Qataris using fifty-five ancestral populations." BMC genetics 13 (2012): 1-10.

**WINPOP**

Paşaniuc, Bogdan, et al. "Inference of locus-specific ancestry in closely related populations." Bioinformatics 25.12 (2009): i213-i221.

**HAPMIX**

Price, Alkes L., et al. "Sensitive detection of chromosomal segments of distinct ancestry in admixed populations." PLoS genetics 5.6 (2009): e1000519.

**ADMIXTURE**

Alexander, David H., John Novembre, and Kenneth Lange. "Fast model-based estimation of ancestry in unrelated individuals." Genome research 19.9 (2009): 1655-1664.

**LAMP**

Sankararaman S, Sridhar S, Kimmel G, Halperin E. Estimating local ancestry in admixed populations. Am J Hum Genet. 2008 Feb;82(2):290-303. PMID: 18252211

**SWITCH**

Sankararaman S, Kimmel G, Halperin E, Jordan MI. On the inference of ancestries in admixed populations. Genome Res. 2008 Apr;18(4):668-75. PMID: 18353809

**HAPAA**

Andreas Sundquist , Eugene Fratkin, Chuong B Do, Serafim Batzoglou. Effect of genetic divergence in identifying ancestral origin using HAPAA. Genome Res. 2008 Apr;18(4):676-82. PMID: 18353807

**SABER**

Tang H, Coram M, Wang P, Zhu X, Risch N. Reconstructing genetic ancestry blocks in admixed individuals. Am J Hum Genet. 2006 Jul;79(1):1-12. PMID: 16773560

**Frappe**

Hua Tang, Jie Peng, Pei Wang, Neil J Risch. Estimation of individual admixture: analytical and study design considerations. Genet Epidemiol. 2005 May;28(4):289-301. PMID: 15712363

**AncestryMAP**

Patterson N, Hattangadi N, Lane B, Lohmueller KE, Hafler DA, Oksenberg JR, Hauser SL, Smith MW, O'Brien SJ, Altshuler D, Daly MJ, Reich D. Methods for high-density admixture mapping of disease genes. Am J Hum Genet. 2004 May;74(5):979-1000. PMID: 15088269

**AdmixMap**

Hoggart CJ, Shriver MD, Kittles RA, Clayton DG, McKeigue PM. Design and analysis of admixture mapping studies. Am J Hum Genet. 2004 May;74(5):965-78. PMID: 15088268

**MALDSOFT**

Giovanni Montana , Jonathan K Pritchard. Statistical tests for admixture mapping with case-control and cases-only data. Am J Hum Genet. 2004 Nov;75(5):771-89. PMID: 15386213

**Figure S1. The examples of the image outputs from AncestryHub using the WGS mode**

Six individuals in two trios (NA12891, NA12892, NA12878, NA19700, NA19701, NA19702) are shown in this figure. The population reference panels were obtained from The 1000 Genomes Project (KGP). AFR, five African populations; SAS, five South Asian populations; EAS, five East Asian populations; EUR, five European populations.

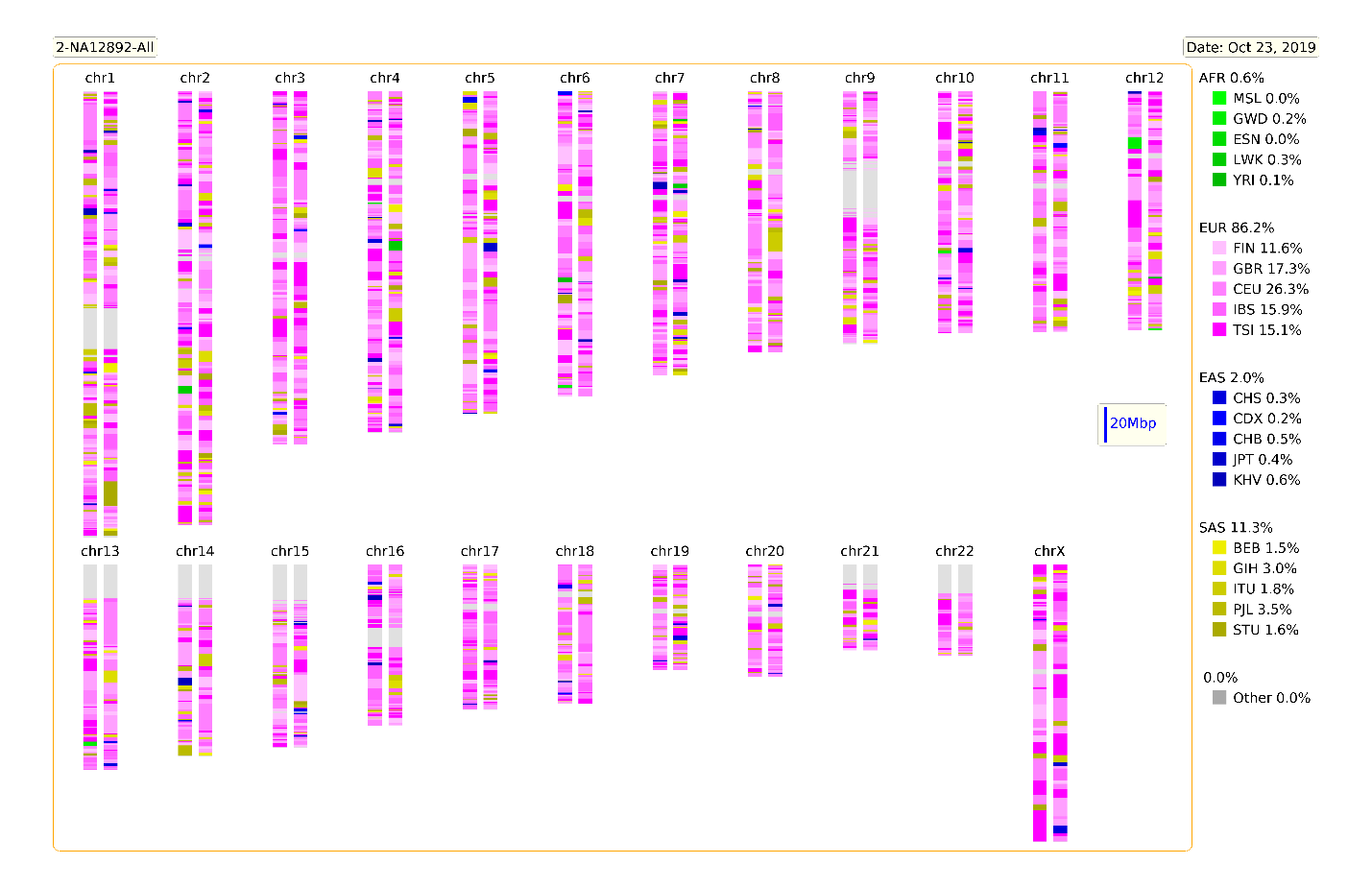

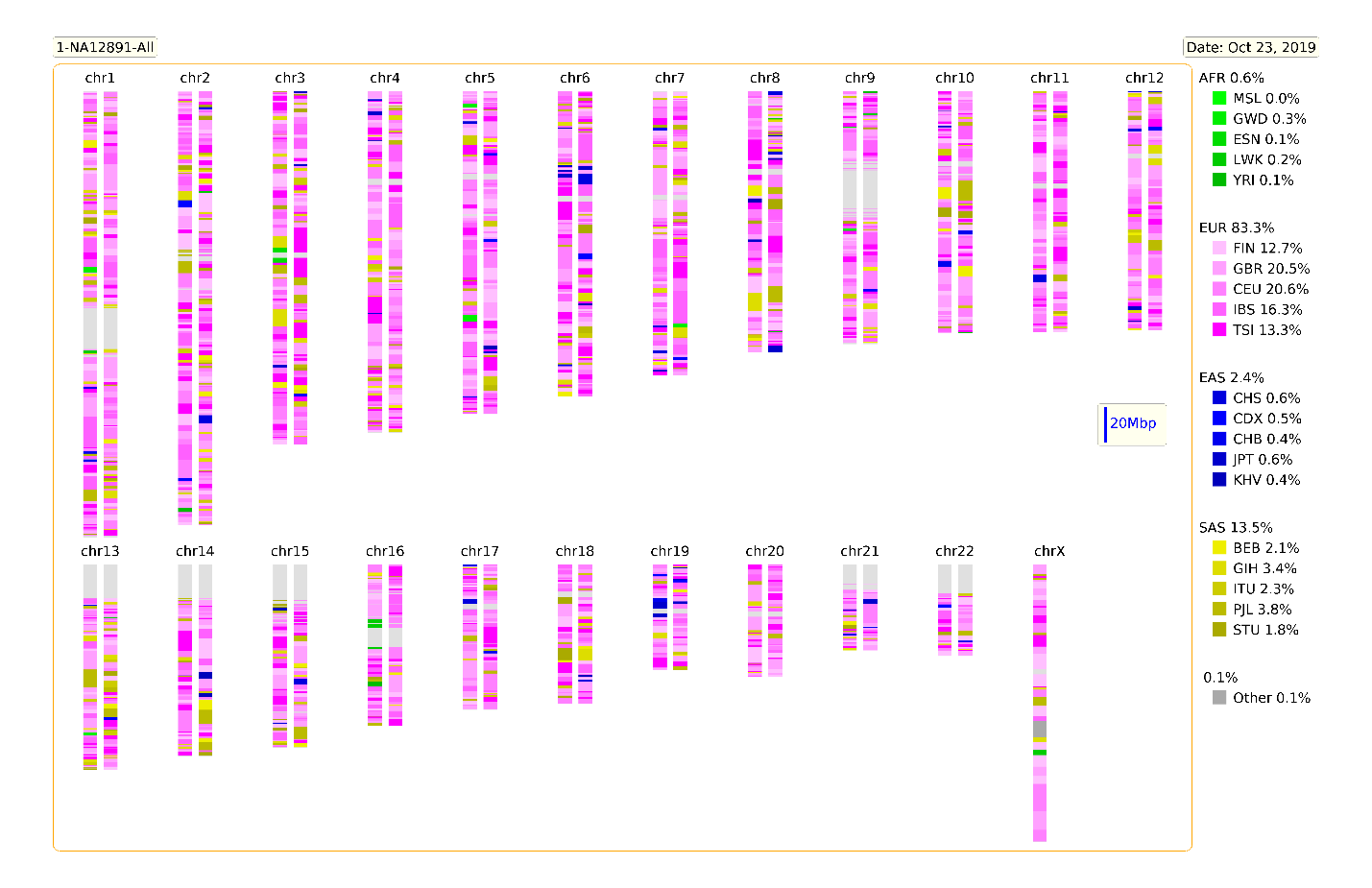

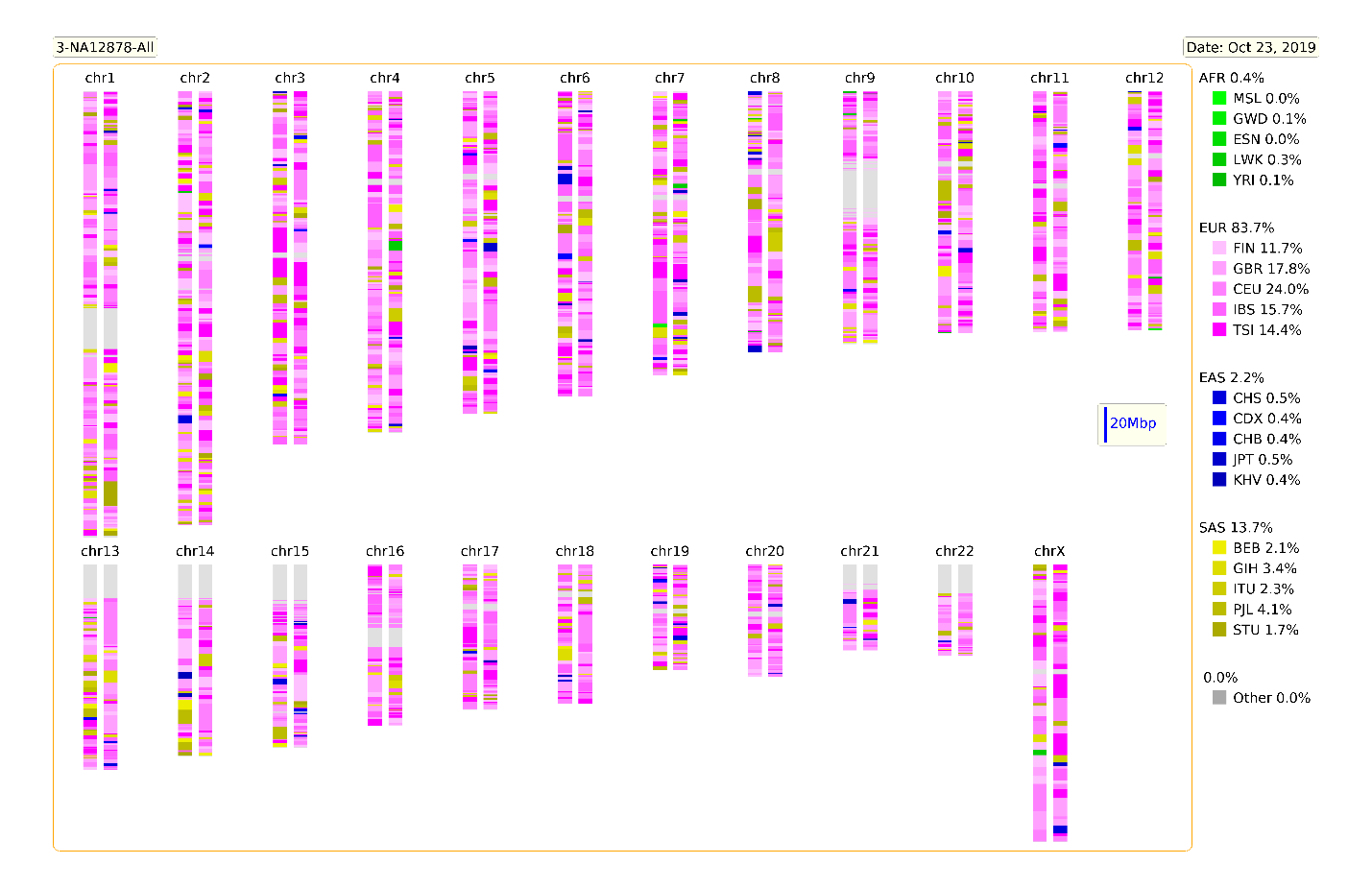

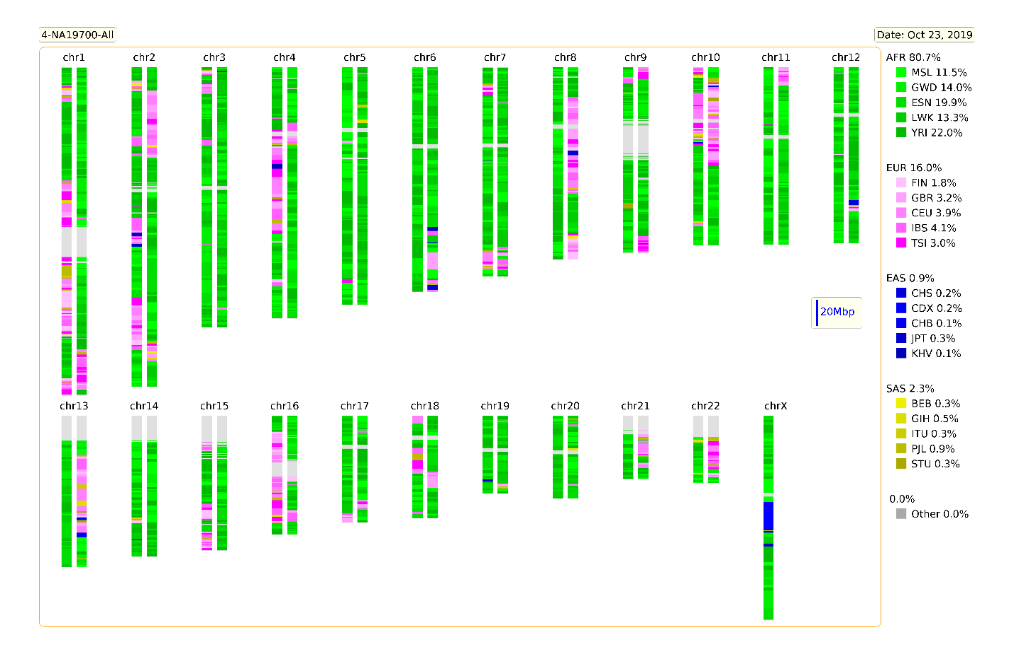

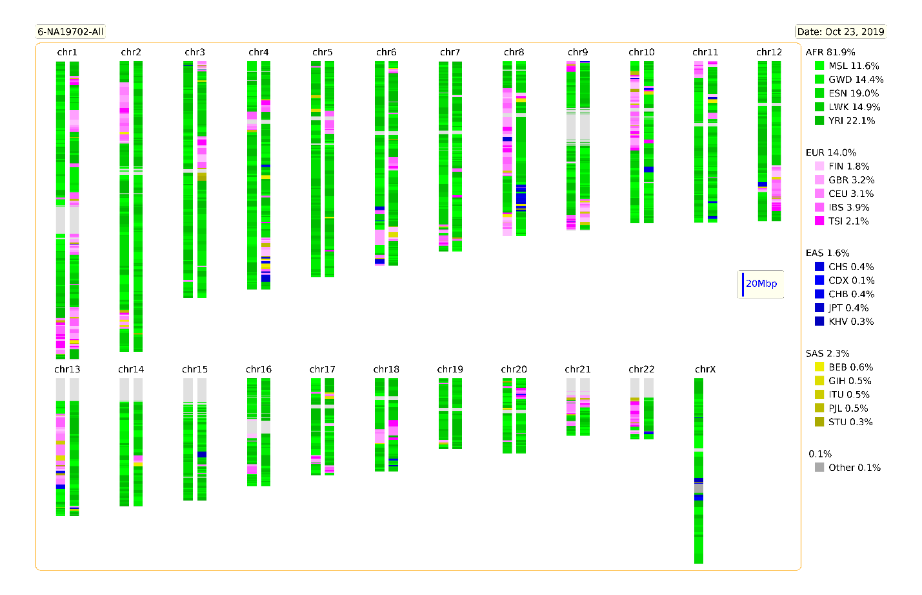

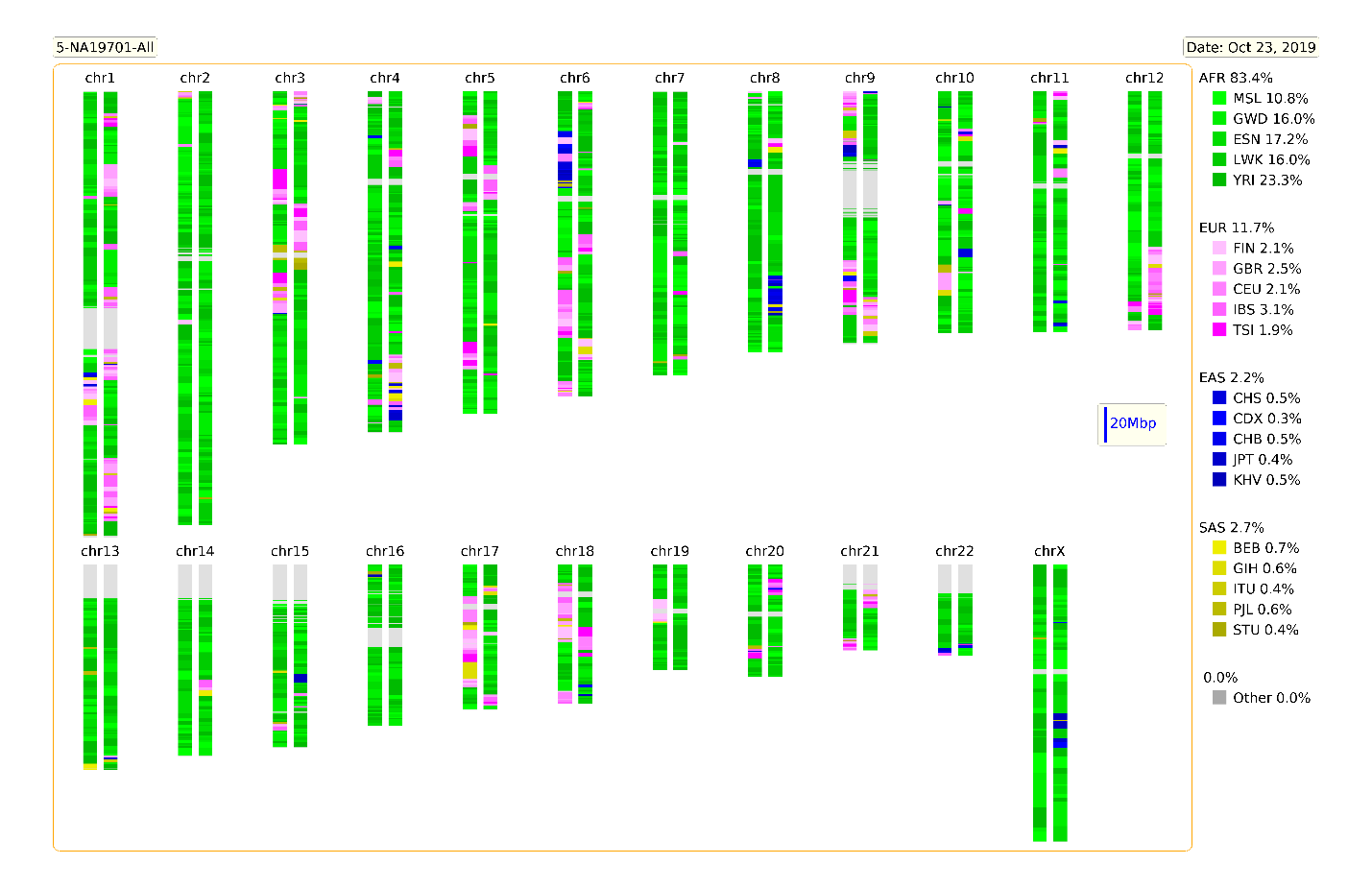

**Figure S2. The examples of the image outputs from AncestryHub using the SCA mode**

The image output of six individuals (NA12891, NA12892, NA12878, NA19700, NA19701, NA19702) on a specific target chromosome region (Chr2:60,000,000-120,000,000) ae shown. The population reference panels were obtained from The 1000 Genomes Project (KGP). AFR, five African populations; SAS, five South Asian populations; EAS, five East Asian populations; EUR, five European populations.

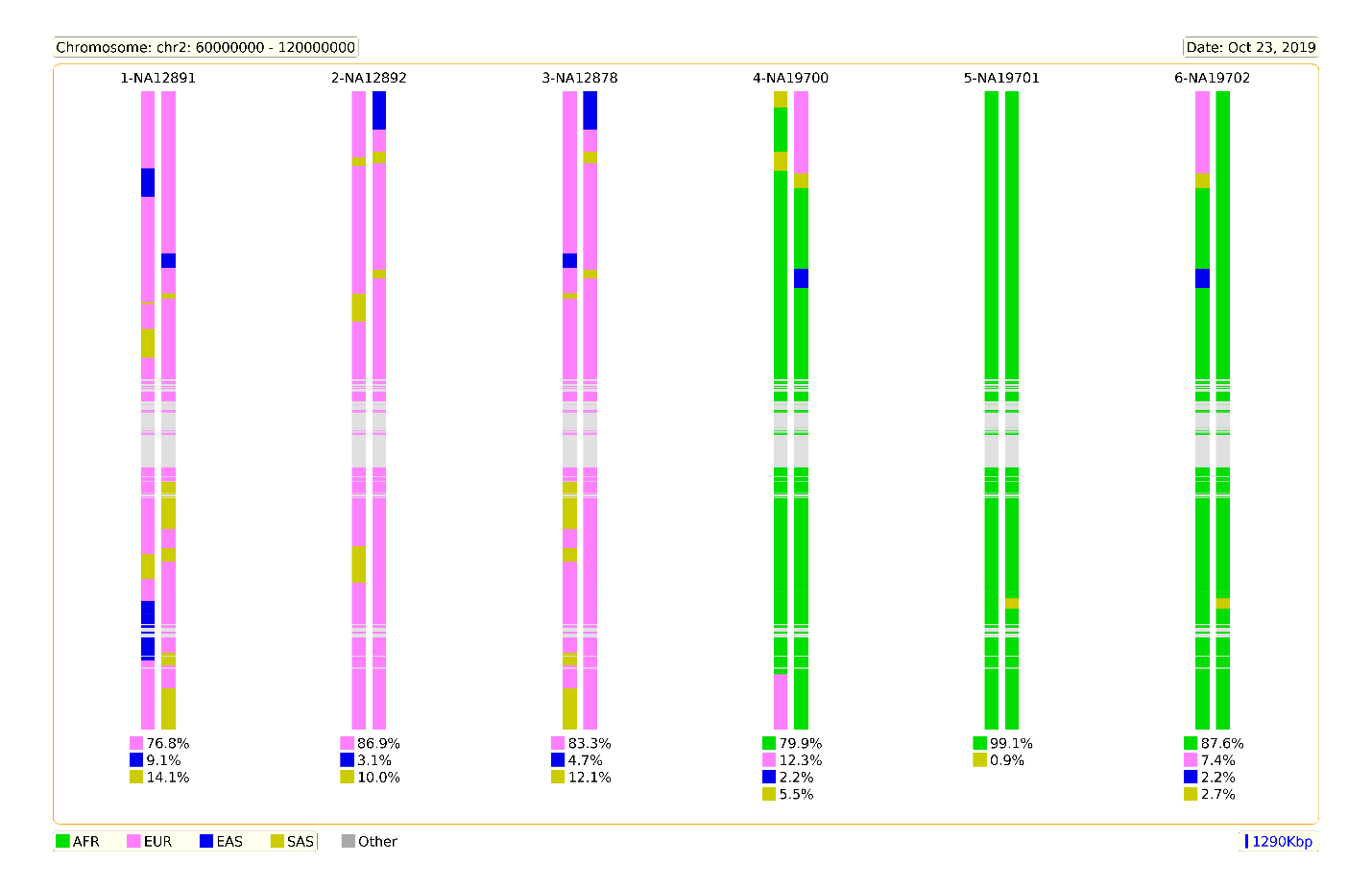

**Figure S3. The examples of the digital outputs from AncestryHub using the WGS mode**

The digital output of AncestryHub WGA results of 6 individuals in two trios (NA12891, NA12892, NA12878, NA19700, NA19701, NA19702) are shown. The population reference panels were obtained from The 1000 Genomes Project (KGP). AFR, five African populations; SAS, five South Asian populations; EAS, five East Asian populations; EUR, five European populations.

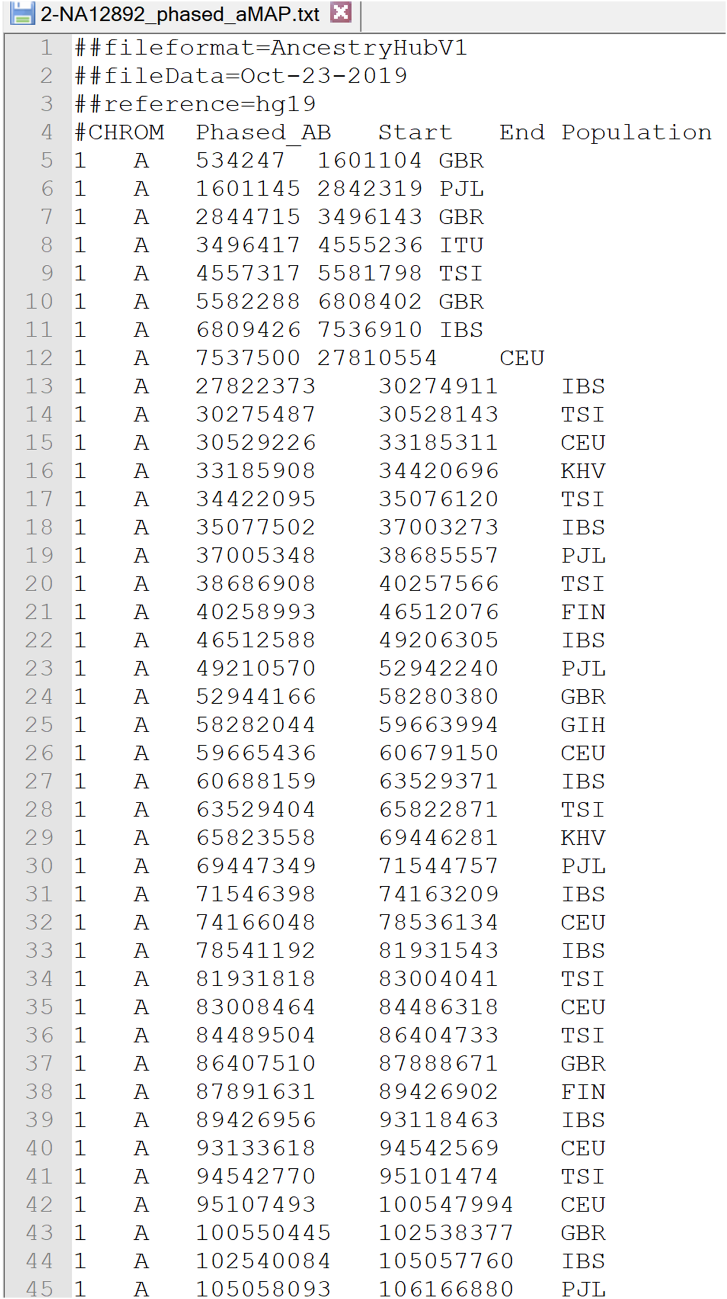

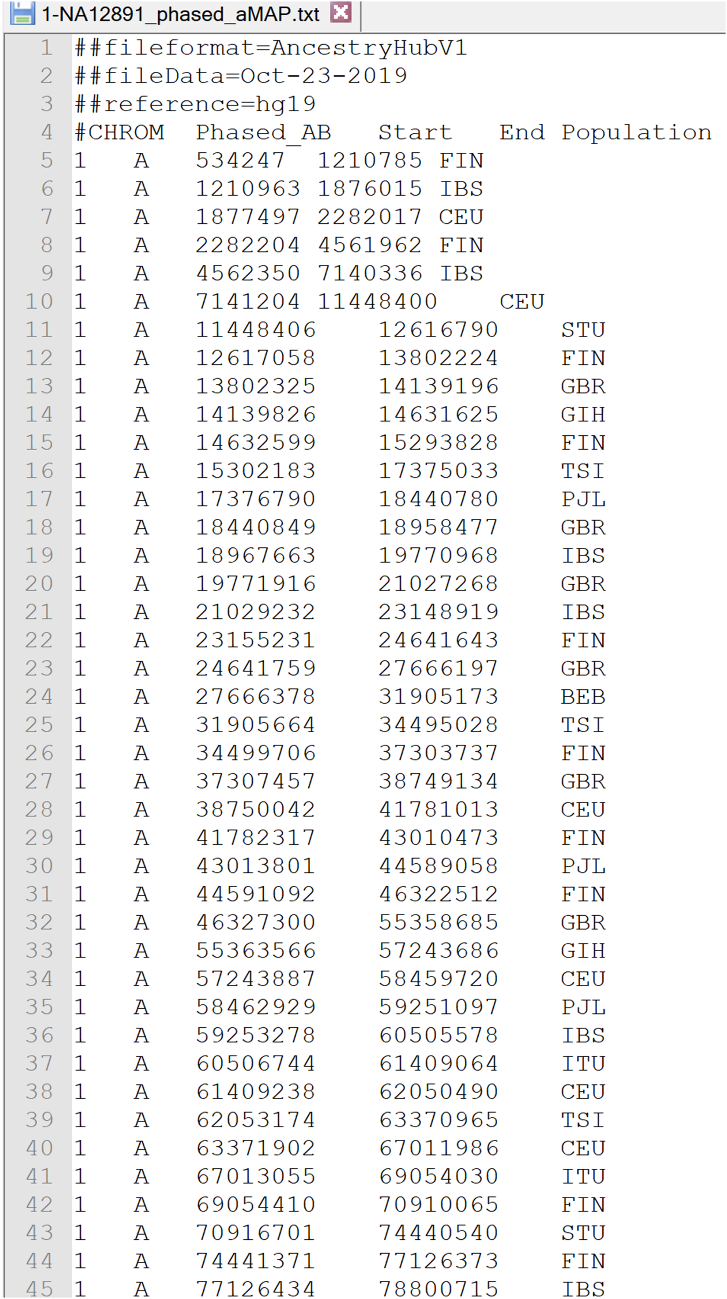

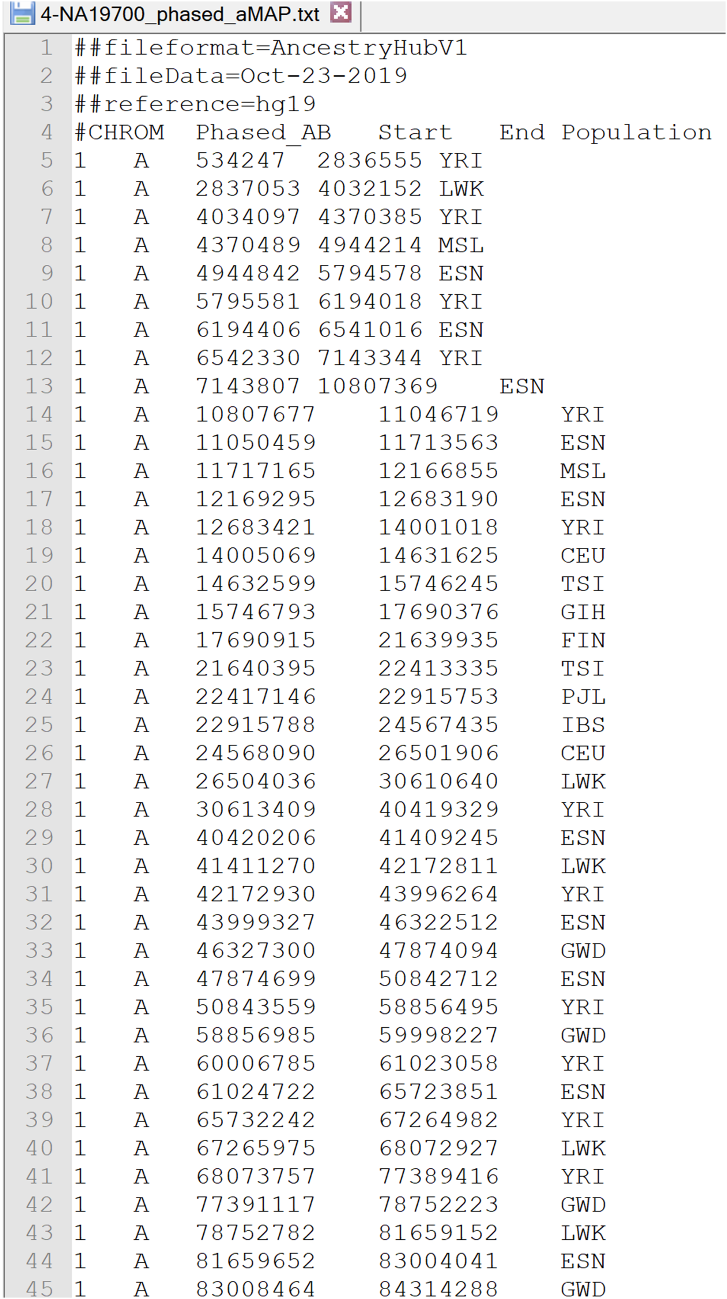

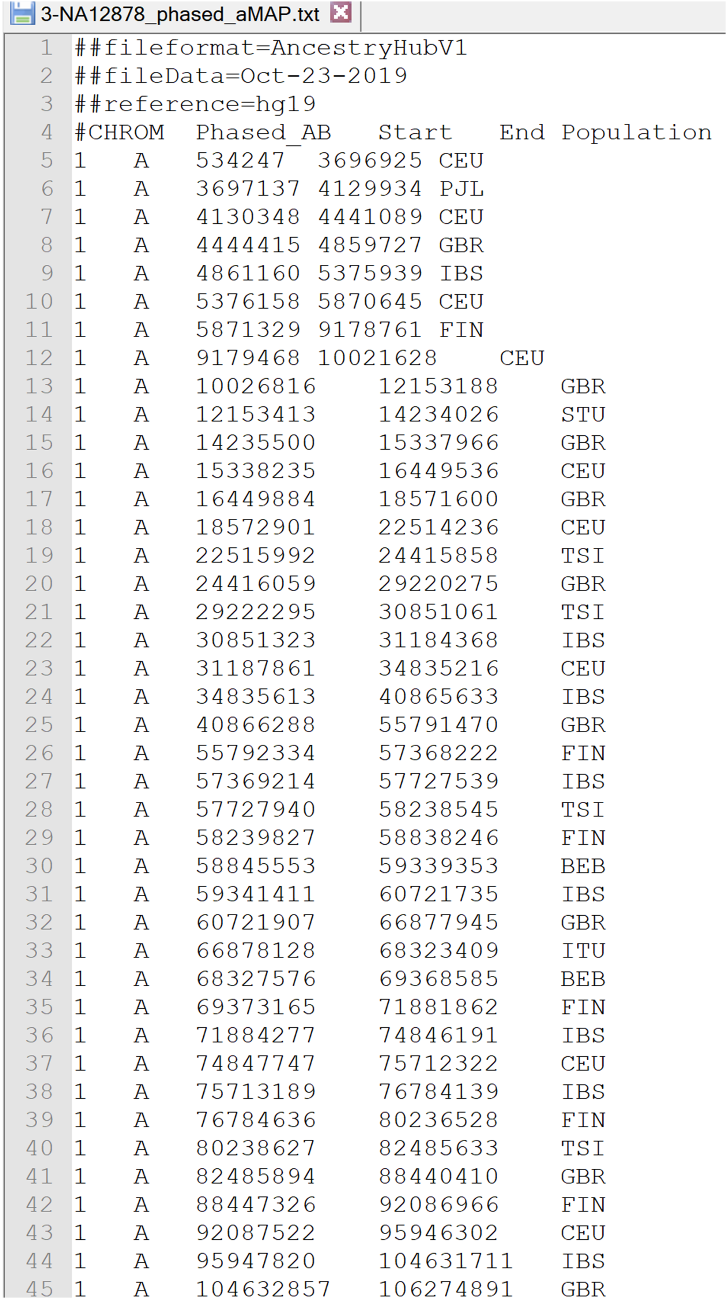

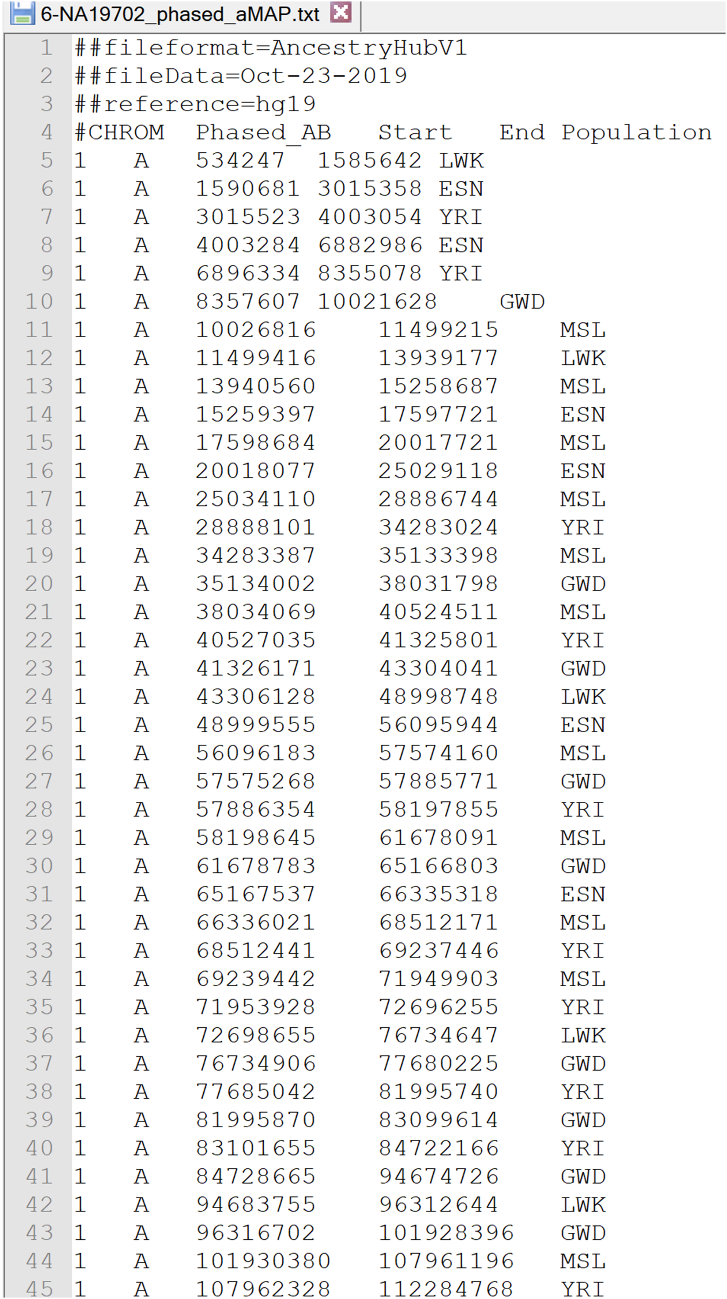

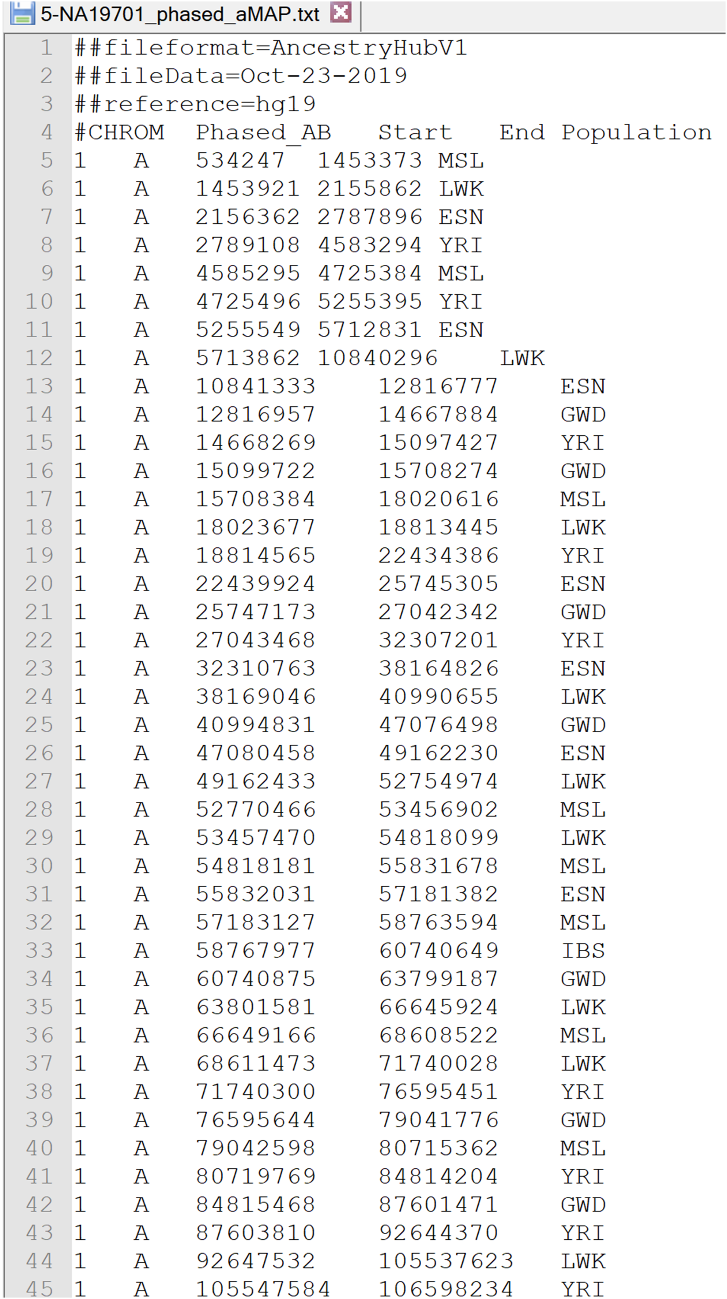

**Figure S4. The examples of the digital outputs from AncestryHub using the SCA mode**

The digital output of the AncestryHub SCA results of 6 individuals in two trios (NA12891, NA12892, NA12878, NA19700, NA19701, NA19702) are shown. The population reference panels were obtained from The 1000 Genomes Project (KGP). AFR, five African populations; SAS, five South Asian populations; EAS, five East Asian populations; EUR, five European populations.

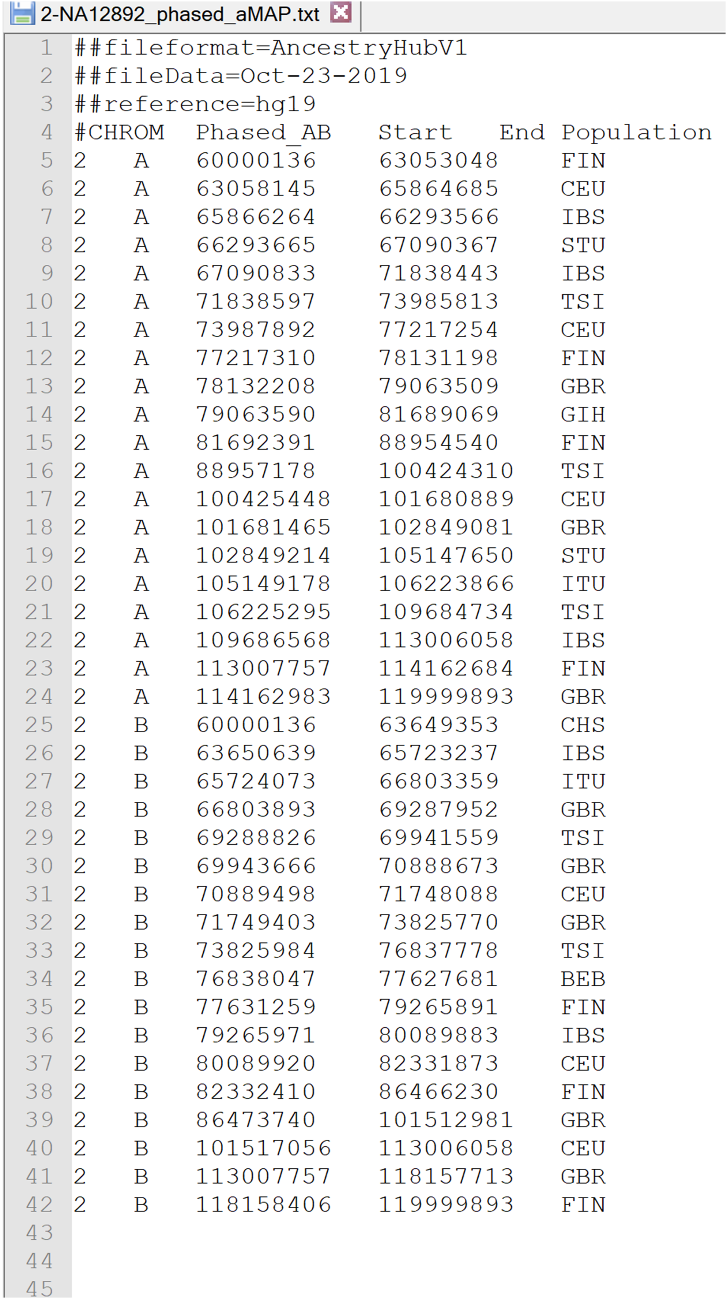

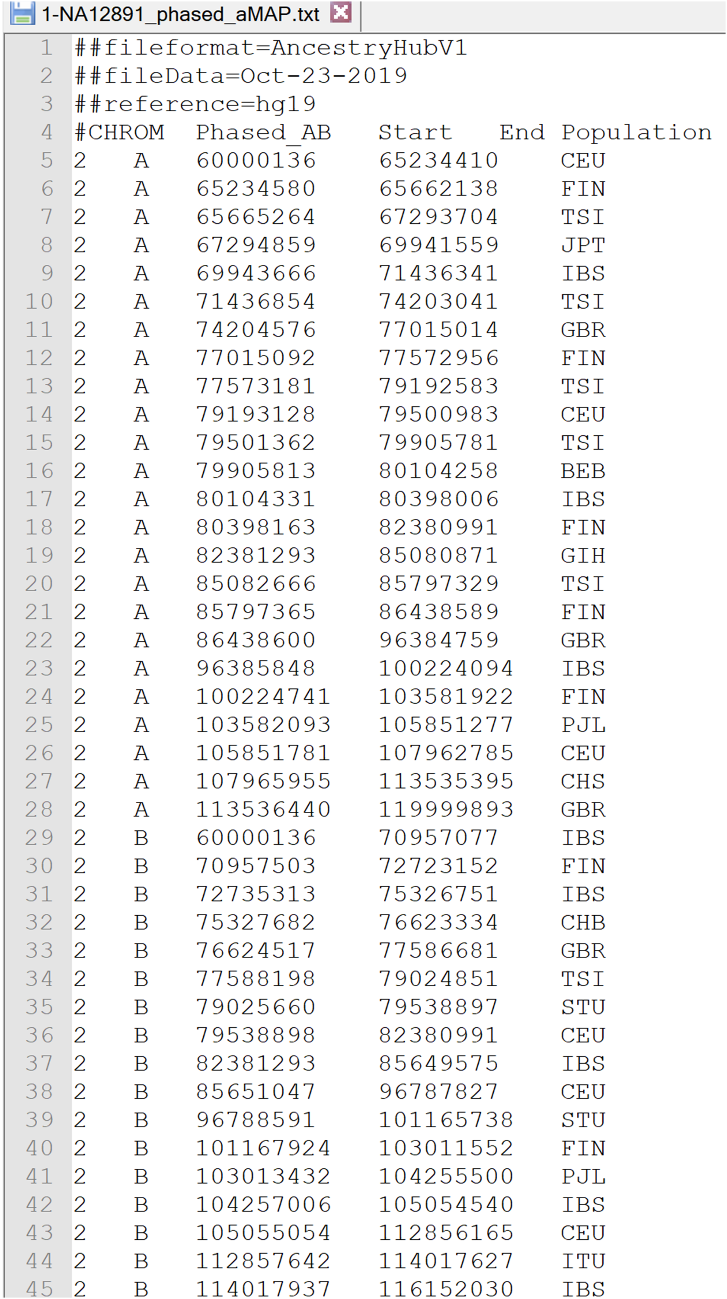

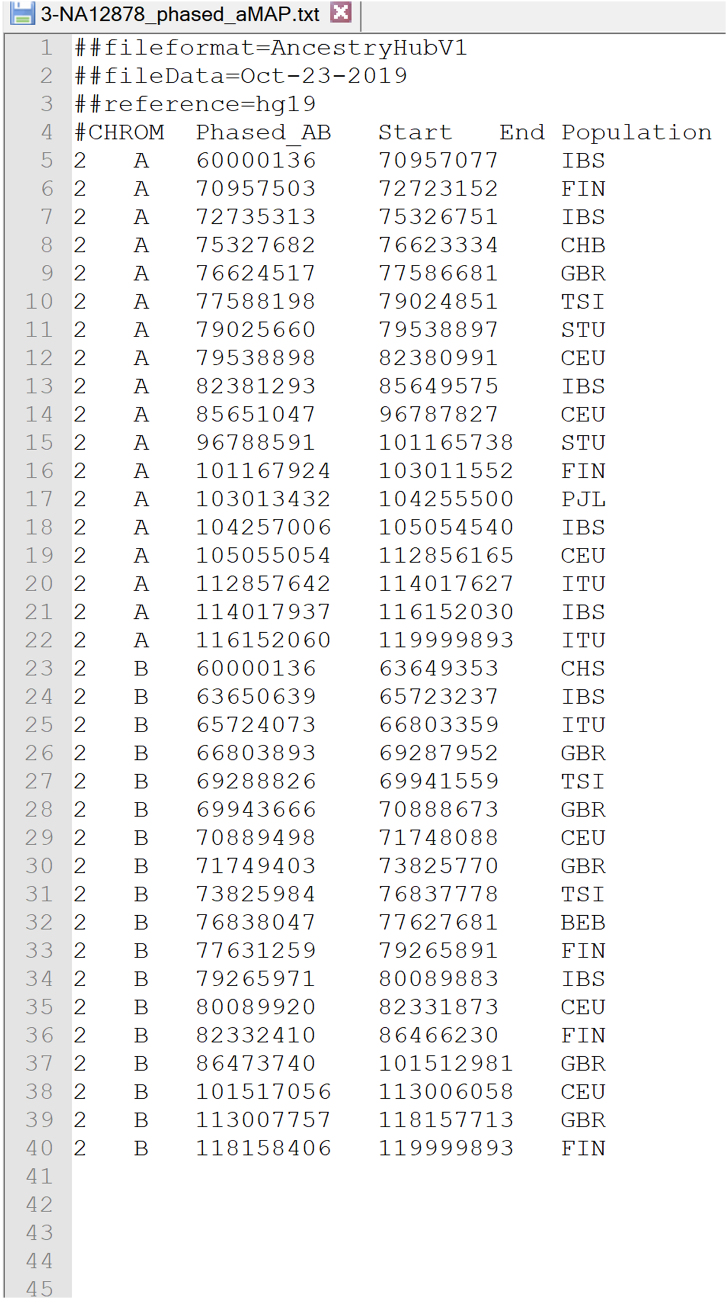

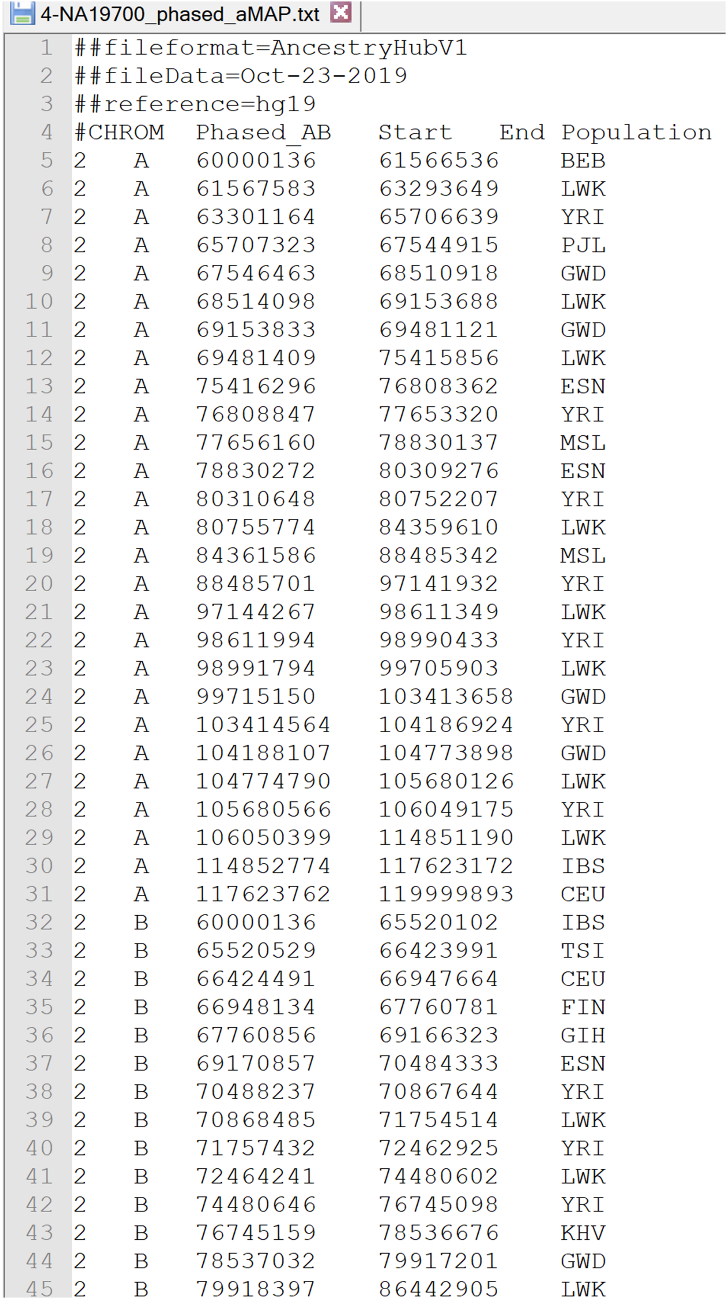

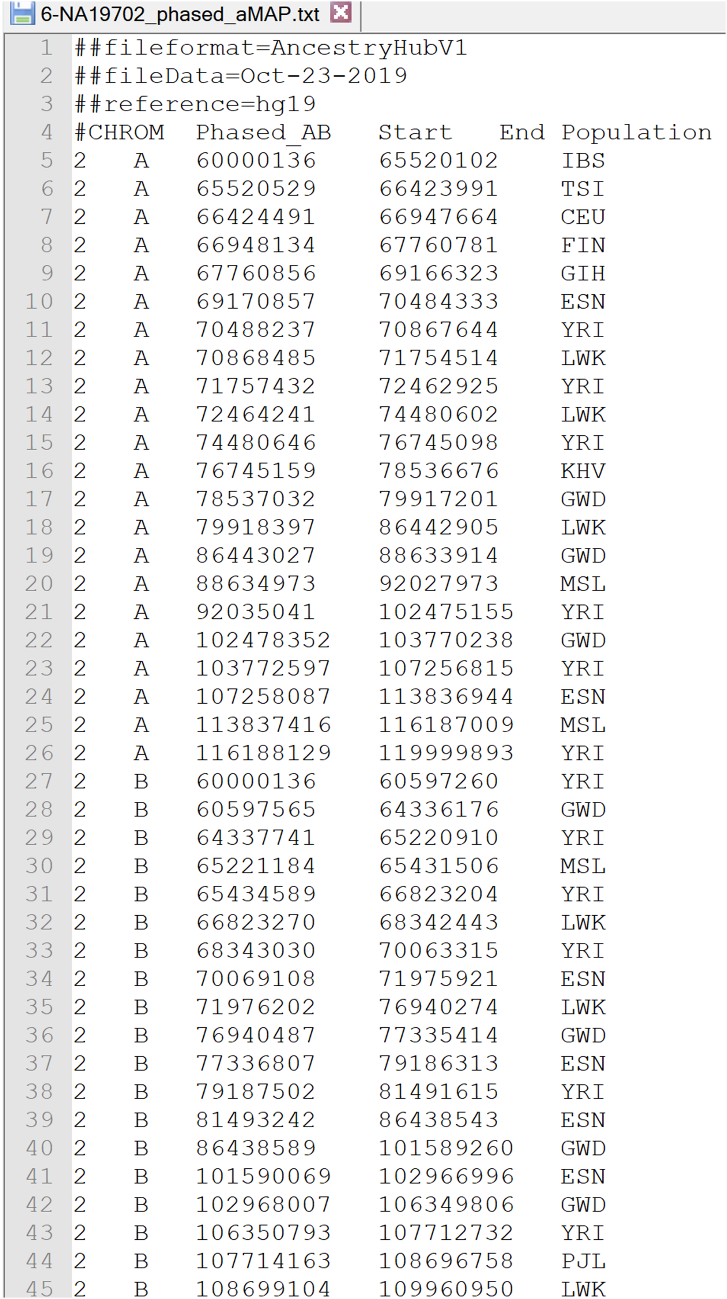

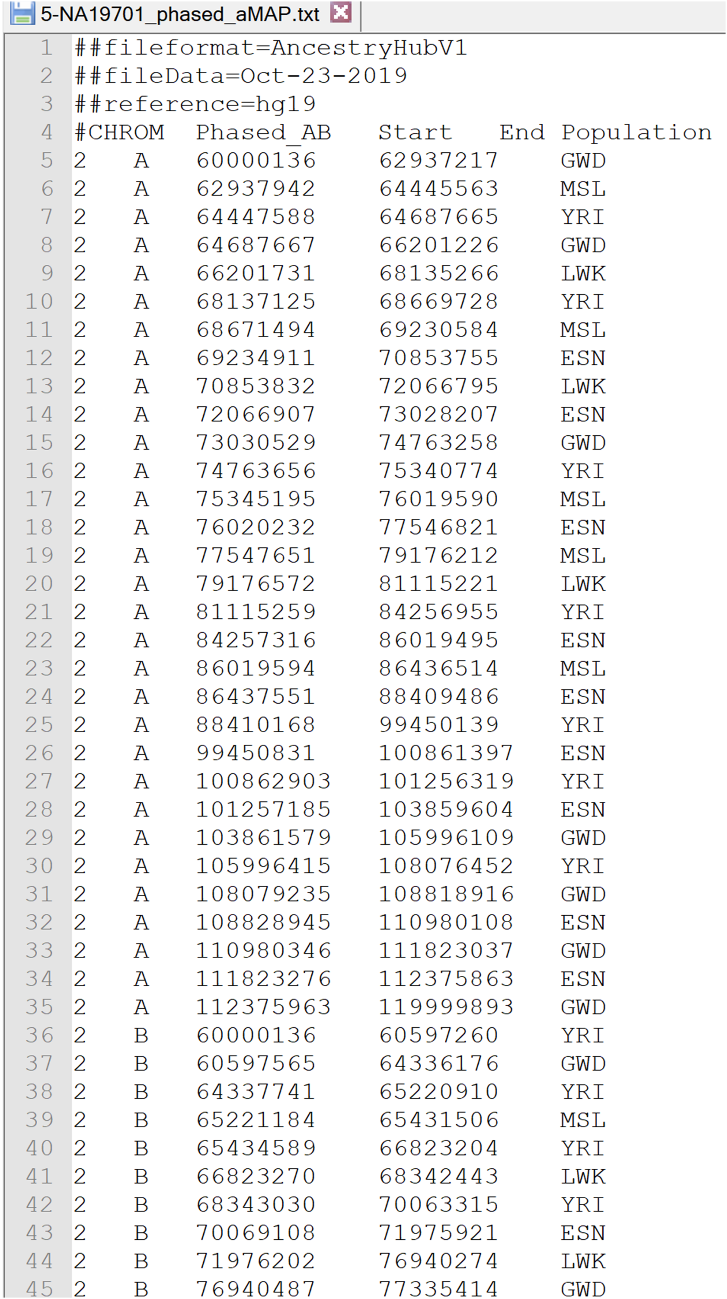

**Table S1. An example of the summary output from AncestryHub using the WGS mode**

| Case ID | 1 | 2 | 3 | 4 | 5 | 6 |
| --- | --- | --- | --- | --- | --- | --- |
| Sample ID | NA12891 | NA12892 | NA12878 | NA19700 | NA19701 | NA19702 |
| MSL | 0.00% | 0.00% | 0.00% | 11.53% | 10.77% | 11.63% |
| GWD | 0.28% | 0.25% | 0.05% | 14.01% | 16.04% | 14.38% |
| ESN | 0.06% | 0.00% | 0.00% | 19.89% | 17.24% | 18.97% |
| LWK | 0.16% | 0.27% | 0.30% | 13.26% | 16.01% | 14.86% |
| YRI | 0.13% | 0.07% | 0.06% | 22.04% | 23.32% | 22.11% |
| FIN | 12.67% | 11.57% | 11.74% | 1.77% | 2.15% | 1.76% |
| GBR | 20.52% | 17.27% | 17.82% | 3.23% | 2.46% | 3.22% |
| CEU | 20.55% | 26.27% | 23.99% | 3.89% | 2.10% | 3.11% |
| IBS | 16.29% | 15.94% | 15.73% | 4.13% | 3.08% | 3.86% |
| TSI | 13.29% | 15.11% | 14.45% | 3.02% | 1.93% | 2.09% |
| CHS | 0.64% | 0.30% | 0.50% | 0.19% | 0.53% | 0.43% |
| CDX | 0.45% | 0.21% | 0.37% | 0.23% | 0.32% | 0.10% |
| CHB | 0.41% | 0.48% | 0.36% | 0.12% | 0.54% | 0.39% |
| JPT | 0.55% | 0.40% | 0.52% | 0.26% | 0.36% | 0.41% |
| KHV | 0.39% | 0.60% | 0.43% | 0.11% | 0.49% | 0.26% |
| BEB | 2.13% | 1.45% | 2.15% | 0.32% | 0.66% | 0.56% |
| GIH | 3.41% | 2.98% | 3.38% | 0.50% | 0.57% | 0.49% |
| ITU | 2.33% | 1.76% | 2.34% | 0.28% | 0.41% | 0.46% |
| PJL | 3.77% | 3.48% | 4.12% | 0.95% | 0.59% | 0.50% |
| STU | 1.85% | 1.59% | 1.71% | 0.28% | 0.43% | 0.30% |
| other | 0.13% | 0.00% | 0.00% | 0.00% | 0.00% | 0.11% |

**Table S2. An example of the summary output from AncestryHub using the SCA mode.**

| Case ID | 1 | 2 | 3 | 4 | 5 | 6 |
| --- | --- | --- | --- | --- | --- | --- |
| Sample ID | NA12891 | NA12892 | NA12878 | NA19700 | NA19701 | NA19702 |
| MSL | 0.00% | 0.00% | 0.00% | 6.64% | 4.83% | 2.76% |
| GWD | 0.00% | 0.00% | 0.00% | 10.27% | 30.28% | 18.14% |
| ESN | 0.00% | 0.00% | 0.00% | 10.62% | 26.00% | 19.91% |
| LWK | 0.00% | 0.00% | 0.00% | 29.74% | 15.11% | 17.05% |
| YRI | 0.00% | 0.00% | 0.00% | 22.66% | 22.91% | 29.76% |
| FIN | 11.73% | 17.82% | 11.72% | 0.95% | 0.00% | 0.95% |
| GBR | 12.57% | 23.21% | 17.19% | 0.00% | 0.00% | 0.00% |
| CEU | 20.44% | 21.84% | 26.82% | 3.08% | 0.00% | 0.54% |
| IBS | 23.22% | 11.93% | 21.97% | 7.18% | 0.00% | 4.82% |
| TSI | 8.85% | 12.09% | 5.55% | 1.14% | 0.00% | 1.14% |
| CHS | 4.09% | 3.06% | 3.06% | 0.00% | 0.00% | 0.00% |
| CDX | 0.00% | 0.00% | 0.00% | 0.00% | 0.00% | 0.00% |
| CHB | 1.61% | 0.00% | 1.61% | 0.00% | 0.00% | 0.00% |
| JPT | 3.36% | 0.00% | 0.00% | 0.00% | 0.00% | 0.00% |
| KHV | 0.00% | 0.00% | 0.00% | 2.21% | 0.00% | 2.21% |
| BEB | 0.37% | 1.01% | 1.01% | 1.47% | 0.00% | 0.00% |
| GIH | 2.00% | 3.54% | 0.00% | 1.85% | 0.00% | 1.85% |
| ITU | 4.98% | 2.58% | 6.24% | 0.00% | 0.00% | 0.00% |
| PJL | 3.02% | 0.00% | 1.07% | 2.19% | 0.88% | 0.88% |
| STU | 3.76% | 2.93% | 3.76% | 0.00% | 0.00% | 0.00% |
| other | 0.00% | 0.00% | 0.00% | 0.00% | 0.00% | 0.00% |

**Table S3. The population Reference Panels from the 1,000 Genomes Project.**

| Population Code | Population Description | Super  Population  Code | The number of  haplotypes |
| --- | --- | --- | --- |
| ACB | African Caribbeans in Barbados | AFR | 192 |
| ASW | Americans of African Ancestry in SW USA | AFR | 122 |
| ESN | Esan in Nigeria | AFR | 198 |
| GWD | Gambian in Western Divisions in the Gambia | AFR | 226 |
| LWK | Luhya in Webuye, Kenya | AFR | 198 |
| MSL | Mende in Sierra Leone | AFR | 170 |
| YRI | Yoruba in Ibadan, Nigeria | AFR | 216 |
| CLM | Colombians from Medellin, Colombia | AMR | 188 |
| MXL | Mexican Ancestry from Los Angeles USA | AMR | 128 |
| PEL | Peruvians from Lima, Peru | AMR | 170 |
| PUR | Puerto Ricans from Puerto Rico | AMR | 208 |
| CDX | Chinese Dai in Xishuangbanna, China | EAS | 186 |
| CHB | Han Chinese in Beijing, China | EAS | 206 |
| CHS | Southern Han Chinese | EAS | 210 |
| JPT | Japanese in Tokyo, Japan | EAS | 208 |
| KHV | Kinh in Ho Chi Minh City, Vietnam | EAS | 198 |
| CEU | Utah Residents (CEPH) with Northern and Western European Ancestry | EUR | 198 |
| FIN | Finnish in Finland | EUR | 198 |
| GBR | British in England and Scotland | EUR | 182 |
| IBS | Iberian Population in Spain | EUR | 214 |
| TSI | Toscani in Italia | EUR | 214 |
| BEB | Bengali from Bangladesh | SAS | 172 |
| GIH | Gujarati Indian from Houston, Texas | SAS | 206 |
| ITU | Indian Telugu from the UK | SAS | 204 |
| PJL | Punjabi from Lahore, Pakistan | SAS | 192 |
| STU | Sri Lankan Tamil from the UK | SAS | 204 |

Because six of these populations (ACB, ASW, CLM, MXL, PEL, PUR) belong to heavily admixed populations ancestrally, we exclude these 6 KGP populations in our built-in reference panels within the AncestryHub.
